# Two gut microbes are necessary and sufficient for normal cognition in *Drosophila melanogaster*

**DOI:** 10.1101/593723

**Authors:** Michael DeNieu, Kristin Mounts, Mollie Manier

## Abstract

It is widely accepted that the gut microbiome can affect various aspects of brain function, including anxiety, depression, learning, and memory. However, we know little about how individual microbial species contribute to communication along the gut-brain axis. Vertebrate microbiomes are comprised of hundreds of species, making it difficult to systematically target individual microbes and their interactions. Here, we use *Drosophila melanogaster* as a simple model organism to tease apart individual and combined effects of gut microbes on cognition. We used an aversive phototactic suppression assay to show that two dominant gut commensals in our lab stock, *Lactobacillus* and *Acetobacter*, are necessary and sufficient for normal learning and short-term memory relative to flies with a conventional microbiome. We also demonstrate that microbes did not affect their hosts’ ability to detect the aversive learning stimulus (quinine), suggesting that our results were due to decreased cognition and not sensory deficits. We thus establish *Drosophila* as a model for elucidating mechanisms of gut-brain communication at the level of individual bacterial species.

## Introduction

Within the last decade, there has been an explosion of studies showing that gut microbes affect critical organismal functions, particularly communication along the gut-brain axis [1]. Early studies found a link between the gut microbiota and anxiety [2–4], depression [5], learning [6], and memory [7] suggesting that gut microbes impact more than just their immediate surroundings. Since then, other studies have found that disrupting the gut microbiota through diet [7], pathogens [8], antibiotics [9, 10], or probiotics [11–14] can all affect learning. Moreover, disease models for inflammatory bowel disease [15, 16], diabetes [17], and schizophrenia [18] have associated the loss of homeostasis of the gut community (dysbiosis) with learning and memory deficits in experimental animals. These deficits were ameliorated with probiotic treatment, suggesting that symptoms experienced by patients may be treated with probiotics as well.

If probiotic therapy is to become an effective treatment for dysbiosis of the microbial community, we need a better understanding of how individual community members influence cognition alone and in combination. We will then be able to target specific microbes or species combinations to more effectively address underlying issues in brain function. We know that diversity of the gut microbiota plays a role (some beneficial species have been identified), but it is not fully understood how individual members of the microbiota contribute, because significantly variable communities can nonetheless be healthy [19–21].

We chose to test learning and memory in *Drosophila*, because the microbiota in flies is much simpler than in vertebrates [22], allowing us to generate single and combinatorial microbial associations that comprise the bulk of the total microbiota. In this way we can begin to dissect community function and develop this system to further probe mechanisms of gut-brain axis communication. We isolated two strains of bacteria from our *D. melanogaster* wild type strain and created populations of flies that were axenic, monoassociated with each microbe, biassociated with two microbes, and a conventional control with the entire microbiota. We then used the phototaxis suppression assay to dissect the individual and combined effects of each microbe in a test of learning and memory. Our assay combined both visual and gustatory cues, whereas most previous studies focused primarily on memory using the novel object recognition test [8–10, 13–15, 18], and only a few studies tested a non-visual sensory modality [12, 14]. We found that removing the microbiota decreased performance in both learning and memory, and that association with a single microbe was not sufficient to restore wild type behavior. However, the biassociation of the two microbes was sufficient, suggesting that effects may be additive or interactive.

## Materials and Methods

We began by characterizing the major taxa in the gut microbiomes of our wild type lab population (LHm). Previous work has shown that *D. melanogaster* laboratory strains typically host relatively few OTUs, dominated by *Lactobacillus* and *Acetobacter* [23]. We homogenized approximately 20 whole flies in 200 *µ*L of mMRS broth [24], plating the resulting solution onto 100 mm mMRS plates and incubating at 30°C for 48 hours. We found that *Lactobacillus* and *Acetobacter* also dominated our lab stock, with no other bacteria detectable by our culturing methods. We visually identified and isolated these bacteria by growing individual colonies in 20 mL of mMRS broth for an additional 48 hours and freezing the cultures at −80°C in 50% glycerol and distilled H_2_O. These cultures were used later to generate a panel of gnotobiotic flies.

We then cleared our LHm flies of intracellular symbionts, including *Wolbachia*, to ensure that we could generate truly germ-free populations. To do this, we reared LHm flies on sucrose and yeast medium at room temperature under ambient light conditions with tetracycline antibiotic for eight consecutive generations. The absence of *Wolbachia* was verified by PCR amplification of 16S ribosomal RNA. We then generated germ-free flies by sterilizing eggs in 0.6% hypochlorite solution and transferred approximately 100 eggs each into sterile 50 mL conical tubes in a laminar flow hood. Axenic flies were reared to adulthood without further manipulation. Monoassociation treatments were supplemented with 100 *µ*L of 48-hour liquid media from *Acetobacter* or *Lactobacillus* cultures, and Combined treatment tubes received 50 *µ*L from both cultures in a separate laminar flow hood to avoid contamination. Conventional flies were produced by allowing 25 males from the base stock to inoculate each tube for 18-24 hours before adding eggs. Shortly after eclosion, all gnotobiotic adults were transferred without anesthesia into polyethelene vials with standard media that had been autoclave sterilized and aged 24-48 hours to verify sterility (lack of microbial growth). We thus generated gnotobiotic flies that carried monoassociations with *“Lactobacillus”* or “*Acetobacter*”, a biassociation with both strains (”Combined”), or neither (”Axenic”). These treatments were compared with “Conventional” flies hosting the full gut microbiome. Because identifying individual bacterial species by culturing is a limited approach, we expect that Conventional flies will carry additional unidentified gut microbes. All flies were aged 3-5 days post-eclosion when assayed.

We used the aversive phototactic suppression assay [25] to assess learning and memory of individual flies. Individual testing eliminated the potential effect of social influence on decision-based behaviors [26]. This assay tests the fly’s ability to associate light with a bitter substance (quinine) and suppress its innate positive phototaxis. To do this, we used a T-maze with a central gate separating a foil-covered dark chamber and a light chamber with the bulb of a goose-neck dissection lamp placed at the end. A 3 cm strip of filter paper was placed into the light chamber near the light source as a vehicle for liquid stimuli that the fly would encounter when moving toward the light. For a benign stimulus, we used 400 *µ*L distilled water, and for the aversive stimulus we used 400 *µ*L of 100 mM quinine. We used separate, dedicated chambers for each stimulus to avoid cross contamination. For each trial, a single fly was aspirated into the dark chamber to acclimate for 30 seconds, after which the light was turned on, and the gate lifted. Flies that entered the light chamber within 30 seconds were scored as phototactic, and flies that remained in the dark chamber were scored as non-phototactic. Once the fly was scored, it was tapped back into the dark chamber for another 30-second acclimation period, and the process was repeated for a total of 16 trials. Flies entering the light chamber generally walked all the way toward the light and encountered the filter paper. All assays were performed in a dark room at 23°C under dim red light.

Flies from each microbiota treatment were subjected to one of three assays. The *phototaxis control assay* consisted of 16 trials with water only in order to establish a baseline phototaxis response for each gnotobiotic treatment. In the *learning assay*, flies were exposed to quinine for all 16 trials to assess their ability to associate quinine with the light. In the *memory assay*, flies were exposed to the quinine for the first eight trials and water for the final eight trials to determine if they could remember the initial association and continue to avoid the light. For all assays, a phototactic response was scored as a binary variable (yes/no), with a positive response if the fly entered the light chamber. For all assays, the last eight trials represented the testing phase, in which we counted the number of phototactic responses. All flies were pre-screened for positive phototaxis in a separate, empty light chamber before each assay. Trials took place between 8 am and 6 pm in a dark room. Because of the large time range, experiments were performed in 30 blocks fully randomized by microbiota treatment and assays type.

For statistical analysis, we used the total number of phototactic trials during the testing phase as the response variable in a linear mixed effects analysis using the lmer function in lme4 v1.1-20 [27] in R v5.2.3 [28]. We fit a model with microbiota treatment (Conventional, Axenic, *Lactobacillus, Acetobacter*, or Combined), assay treatment (phototaxis, learning, or memory) and their interaction as fixed effects, with the date each block was performed as a random effect. All estimates report the mean, standard error. The p-values for model fixed effects were generated using a parametric bootstrap in the bootMer function in lme4 with 100000 simulations.

To ensure that the results of the learning assay were due to differences in cognition rather than taste perception, we tested each gnotobiotic treatment’s gustatory response to quinine in colored agar. To do this, we used three different taste assays: no quinine, quinine in red agar, and quinine in blue agar. The control treatment contained no quinine, in order to assess a color preference. The red or blue quinine treatments had 100 mM quinine in the red or blue 1% agar (with McCormick food coloring), respectively, controlling for any inherent color preference. A solution of live yeast in water was added to the surface of each well to stimulate feeding. We reared flies on our standard sterile media, collected flies 3-5 days old under light CO_2_ anesthesia into vials at densities of 25 flies per vial. After 24 hours acclimation, 50 flies (equal sex ratio) were bumped into a sealed plastic container with a 16-well microtiter plate filled in alternating patterns with red or blue agar. This experiment was repeated for each microbiota and quinine treatment group, such that approximately 100 flies were assayed for each combination. Containers were enclosed in a cardboard box to exclude light and placed in a 23°C incubator for 24 hours. Flies were then anesthetized with CO_2_, and abdomen color recorded as either blue, red, or purple. Because fewer than 6% flies in the control assay were purple, and no flies were purple in either quinine assay, we removed them from the analysis. For statistical analysis, we calculated proportions of flies consuming each color in each gnotobiotic treatment in the control assay. We used those proportions to generate matching expected count values for each gnotobiotic treatment in chi-squared goodness of fit tests for the assays with the quinine in the blue agar and the red agar respectively. Reported p-values were generated using Monte Carlo simulation with 100000 replicates.

## Results

All gnotobiotic treatments showed significant levels of learning, as measured by reductions in phototaxis in the presence of quinine relative to the phototaxis control assay (Figure 1*a*). In the learning assay, control flies had the largest reduction in phototaxis as predicted (Δphototaxis = −3.66 ± 0.55, *p* < 0.0001) However, the Axenic (Δphototaxis = −1.73 ± 0.77, *p* = 0.015), *Lactobacillus* (Δphototaxis = −1.67 ± 0.77, *p* = 0.011) and *Acetobacter* treatments (Δphototaxis = −1.60 ± 0.76, *p* < 0.009) had reduced learning relative to Conventional flies. This result suggests that an intact gut microbiome is required for normal cognition, and neither *Lactobacillus* nor *Acetobacter* alone were sufficient to improve learning above that of Axenic flies. A biassociation of both species in the Combined treatment, however, rescued learning (Δphototaxis =-3.10 ± 0.77; *p* = 0.51), suggesting that both bacterial species are necessary and sufficient for normal cognition.

**Figure 1.**
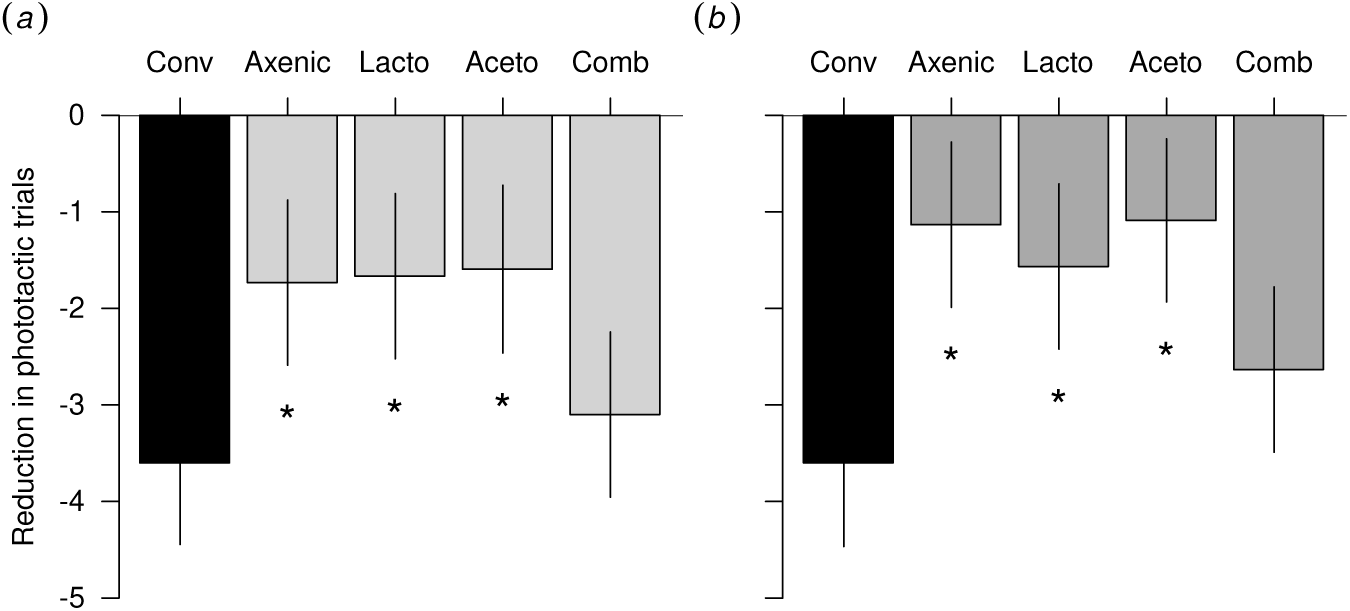
Reduction in phototactic trials during the testing phase of the *(a)* learning (quinine only) and *(b)* memory (quinine, then water) assays relative to the phototaxis control assay (water only), with 95% confidence intervals. Asterisks represent significant differences relative to the Conventional treatment.

A similar outcome was found for the memory assay (Figure 1*b*). All treatments had significant reductions in phototaxis (stats), but the Axenic (Δphototaxis = −1.13 ± 0.77; *p* = 0.001), *Lactobacillus* (Δphototaxis = −1.57 ± 0.77; *p* = 0.008), and *Acetobacter* (Δphototaxis = −1.09 ± 0.77; *p* = 0.0001) treatments performed more poorly relative to Conventional flies. Again, memory was rescued in the Combined treatment (Δphototaxis = −2.63 ± 0.77; *p* = 0.21). Though all treatments were similarly positively phototactic with water (*p*_*microbiota*_ = 0.37), there were minor differences among the microbiota groups, with *Acetobacter* flies showing the largest difference as compared to the conventional flies (Table 1). To account for this, we used this assay as a baseline level of phototaxis specific to each treatment (a relative value of 0 in Figure 1), against which cognition was compared. Thus, minor differences in phototaxis among treatments were accounted for in the learning and memory assessments.

**Table 1.** Average number of phototactic trials in the testing phase for the control, learning, and memory assays for each microbiota treatment, with standard errors in parentheses.

|  | Conv | Axenic | *Lacto* | *Aceto* | Comb |
| --- | --- | --- | --- | --- | --- |
| Control | 5.69 (0.43) | 4.91 (0.54) | 4.82 (0.54) | 4.68 (0.54) | 5.19 (0.54) |
| Learning | 2.09 (0.54) | 3.19 (0.76) | 3.15 (0.76) | 3.09 (0.77) | 2.09 (0.76) |
| Memory | 2.09 (0.55) | 3.79 (0.77) | 3.25 (0.77) | 3.59 (0.77) | 2.55 (0.77) |

In the taste assays, we detected no differences among the gnotobiotic treatments in quinine taste perception. Axenic and *Acetobacter* flies did show a preference for red media, avoiding the blue, while *Lactobacillus*, Combined, and Conventional flies showed no preference (Figure 2*a*). However, all treatments avoided the color containing the quinine, regardless of their base preference (Figure 2*b,c*; *p ≤* 0.01 for all treatments except *Acetobacter* with the blue quinine where avoidance of blue was already strong and only slightly increased). These results suggest that the microbiome treatments were not disrupting taste perception, and that our results reflected differences in cognition.

**Figure 2.**
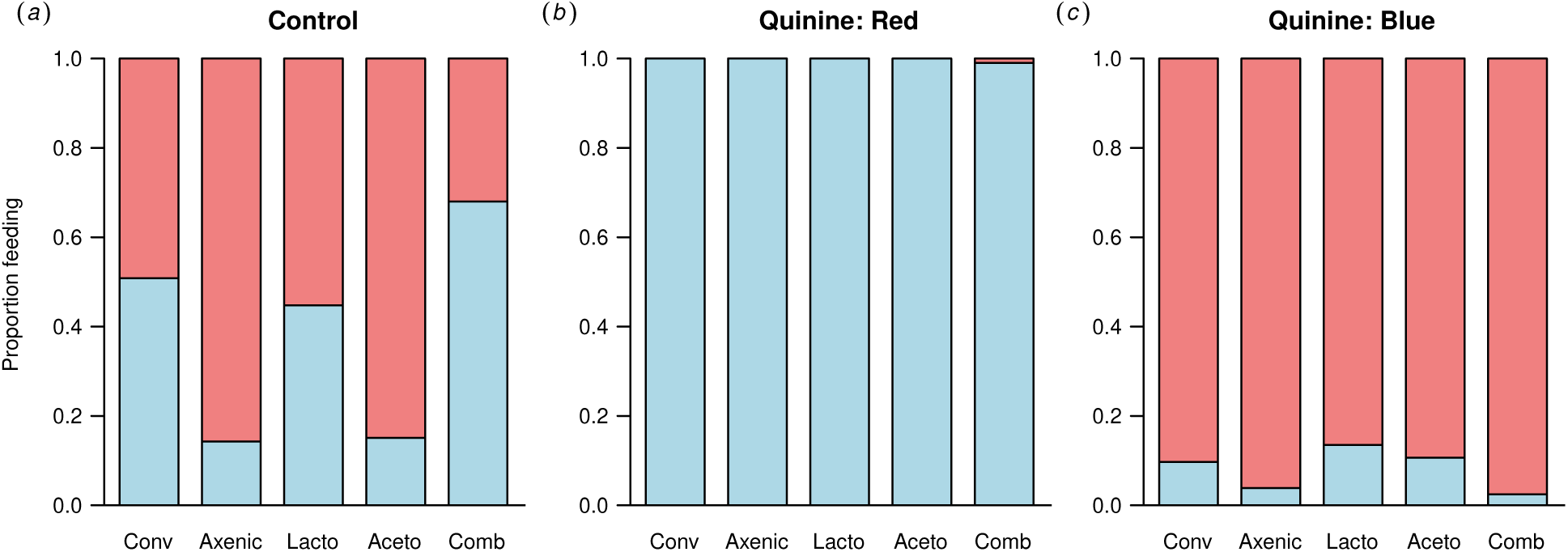
Proportion of flies eating the blue and red agar for *(a)* control with no quinine, *(b)* quinine in red agar only, and *(c)* quinine in blue agar only.

## Discussion

Our findings demonstrate that *Drosophila melanogaster* can be used as a simple model organism for isolating different components of the gut-brain axis, beginning here with individual members of the gut community. We showed that learning deficits can be invoked by manipulating the gut microbiome, and that multiple species are required to rescue normal learning. A significant role of the gut microbiome in learning in both flies and vertebrates suggests that mechanisms of gutbrain communication may be evolutionarily conserved. Indeed, there is widespread homology in neurological mechanisms of learning [29], intestinal structure [30], and overlap in community members between vertebrates and *Drosophila* (e.g., *Lactobacillus* and other Firmicutes; [23, 31]).

The neurological pathways controlling olfaction in *Drosophila* are both more complex [32] and better understood than those for gustation, but there is substantial overlap in how both sensory modalities feed into learning centers in the brain. Signals initiated by bitter taste receptors on the legs are communicated to the brain by type 3 taste projection neurons (TPN3), with axons terminating in the higher brain close to olfactory neural projections [33]. TPN3 neurons directly activate PPL1 dopaminergic neurons [33] that are also required for aversive olfactory learning and target the central complex and mushroom body [34], the site of learning. Indeed, dopamine, serotonin, and GABA all contribute to regulating learning in flies [35–40], with similar functions in vertebrate learning systems [41–43].

These neuroactive peptides can also be produced by gut bacteria themselves. For example, *Lactobacillus brevis*, a common commensal of *D. melanogaster* [23], is a proficient producer of GABA in culture [43], and other *Lactobacillus* species are known to produce dopamine, serotonin, and GABA [44]. Probiotic treatment of mice with *Lactobacillus plantarum* (also commonly found in *D. melanogaster)* increases dopamine and decreases serotonin in the brain and ameliorates anxiety-like behaviors [45]. Additionally, gut commensals have been shown to decrease symptoms of depression by catalyzing the conversion of tryptophan into kynurenine and reducing its availability to produce serotonin [46, 47]. Short-chain fatty acids (SCFAs) are another class of bacterially-derived metabolite that may be associated with brain function [48–51]. In *Drosophila*, the SCFA acetate is produced by *Acetobacter* and regulates host growth and development via the insulin signalling pathway [52–54]. If and how these metabolites directly travel to the central nervous system is unclear, but in mice, acetate produced in the colon can reach and cross the blood-brain barrier to control appetite, accumulating in the hypothalamus and incorporating into GABA within GABAergic neurons [55].

Though the mechanism of action in our study has not yet been elucidated, our results underscore the importance of gut community structure on host phenotype, with emergent properties that exceed the sum of its parts. These results also suggest that probiotic treatments targeting single microbial species may be less effective than treatments targeting the whole community (e.g., fecal transplant). At the same time, it is important to understand the nature of species interactions among gut microbes in order to design the most effective and personalized treatments. Future directions include characterizing metabolites produced by our *Lactobacillus* and *Acetobacter* strains in culture as well as in the guts of mono- and biassociated flies, especially since biogenic amine production by bacteria may be more strain-specific than species-specific [56]. Taken together, we show that the gut microbiota are important for the maintenance of learning and memory in *D. melanogaster*, a simple model organism in which the differences between mono- and biassociations can be easily probed. This approach may prove to be a valuable starting point for understanding the interactions between the microbiota and the host in more complex gut microbiomes.

## References

[1] Gareau MG. 2016 Cognitive function and the microbiome. International review of neurobiology 131, 227–246. (doi:10.1016/bs.irn.2016.08.001).

[2] Rao AV, Bested AC, Beaulne TM, Katzman MA, Iorio C, Berardi JM, Logan AC. 2009 A randomized, double-blind, placebo-controlled pilot study of a probiotic in emotional symptoms of chronic fatigue syndrome. Gut pathogens 1, 6. (doi:10.1186/1757-4749-1-6).

[3] Messaoudi M, Violle N, Bisson JF, Desor D, Javelot H, Rougeot C. 2011 Beneficial psychological effects of a probiotic formulation (lactobacillus helveticus R0052 and bifidobacterium longum r0175) in healthy human volunteers. Gut microbes 2, 4, 256–261. (doi: 10.4161/gmic.2.4.16108).

[4] Messaoudi M, Lalonde R, Violle N, Javelot H, Desor D, Nejdi A, Bisson JF, Rougeot C, Pichelin M, Cazaubiel M, et al.2011 Assessment of psychotropic-like properties of a probiotic formulation (lactobacillus helveticus R0052 and bifidobacterium longum r0175) in rats and human subjects. The British journal of nutrition 105, 05, 755–764. (doi: 10.1017/S0007114510004319).

[5] Steenbergen L, Sellaro R, van Hemert S, Bosch JA, Colzato LS. 2015 A randomized controlled trial to test the effect of multispecies probiotics on cognitive reactivity to sad mood. Brain, behavior, and immunity 48, 258–264. (doi:10.1016/j.bbi.2015.04.003).

[6] Carlson AL, Xia K, Azcarate-Peril MA, Goldman BD, Ahn M, Styner MA, Thompson AL, Geng X, Gilmore JH, Knickmeyer RC. 2018 Infant gut microbiome associated with cognitive development. Biological psychiatry 83, 2, 148–159. (doi:10.1016/j.biopsych.2017.06.021).

[7] Li W, Dowd SE, Scurlock B, Acosta-Martinez V, Lyte M. 2009 Memory and learning behavior in mice is temporally associated with diet-induced alterations in gut bacteria. Physiology & behavior 96, 4–5, 557–567. (doi:10.1016/j.physbeh.2008.12.004).

[8] Gareau MG, Wine E, Rodrigues DM, Cho JH, Whary MT, Philpott DJ, MacQueen G, Sherman PM. 2011 Bacterial infection causes stress-induced memory dysfunction in mice. Gut 60, 3, 307–317. (doi:10.1136/gut.2009.202515).

[9] Ceylani T, Jakubowska-Doğgru E, Gurbanov R, Teker HT, Gozen AG. 2018 The effects of repeated antibiotic administration to juvenile BALB/c mice on the microbiota status and animal behavior at the adult age. Heliyon 4, 6, e00644. (doi:10.1016/j.heliyon.2018.e00644).

[10] Fröhlich EE, Farzi A, Mayerhofer R, Reichmann F, Jacan A, Wagner B, Zinser E, Bordag N, Magnes C, Fröhlich E, et al.2016 Cognitive impairment by antibiotic-induced gut dysbiosis: Analysis of gut microbiota-brain communication. Brain, behavior, and immunity 56, 140– 155. (doi:10.1016/j.bbi.2016.02.020).

[11] Matthews DM, Jenks SM. 2013 Ingestion of mycobacterium vaccae decreases anxiety-related behavior and improves learning in mice. Behavioural processes 96, 27–35. (doi: 10.1016/j.beproc.2013.02.007).

[12] Xiao J, Li S, Sui Y, Wu Q, Li X, Xie B, Zhang M, Sun Z. 2014 Lactobacillus casei-01 facilitates the ameliorative effects of proanthocyanidins extracted from lotus seedpod on learning and memory impairment in scopolamine-induced amnesia mice. PloS one 9, 11, e112773. (doi:10.1371/journal.pone.0112773).

[13] Smith CJ, Emge JR, Berzins K, Lung L, Khamishon R, Shah P, Rodrigues DM, Sousa AJ, Reardon C, Sherman PM, et al.2014 Probiotics normalize the gut-brain-microbiota axis in immunodeficient mice. American journal of physiology. Gastrointestinal and liver physiology 307, 8, G793–802. (doi:10.1152/ajpgi.00238.2014).

[14] Savignac HM, Tramullas M, Kiely B, Dinan TG, Cryan JF. 2015 Bifidobacteria modulate cognitive processes in an anxious mouse strain. Behavioural brain research 287, 59–72. (doi:10.1016/j.bbr.2015.02.044).

[15] Emge JR, Huynh K, Miller EN, Kaur M, Reardon C, Barrett KE, Gareau MG. 2016 Modulation of the microbiota-gut-brain axis by probiotics in a murine model of inflammatory bowel disease. American Journal of Physiology-Gastrointestinal and Liver Physiology 310, 11, G989–G998.

[16] Ohland CL, Kish L, Bell H, Thiesen A, Hotte N, Pankiv E, Madsen KL. 2013 Effects of lactobacillus helveticus on murine behavior are dependent on diet and genotype and correlate with alterations in the gut microbiome. Psychoneuroendocrinology 38, 9, 1738–1747. (doi:10.1016/j.psyneuen.2013.02.008).

[17] Davari S, Talaei SA, Alaei H, Salami M. 2013 Probiotics treatment improves diabetes-induced impairment of synaptic activity and cognitive function: behavioral and electrophysiological proofs for microbiome-gut-brain axis. Neuroscience 240, 287–296. (doi: 10.1016/j.neuroscience.2013.02.055).

[18] Pyndt Jørgensen B, Krych L, Pedersen TB, Plath N, Redrobe JP, Hansen AK, Nielsen DS, Pedersen CS, Larsen C, Sørensen DB. 2015 Investigating the long-term effect of subchronic phencyclidine-treatment on novel object recognition and the association between the gut microbiota and behavior in the animal model of schizophrenia. Physiology & behavior 141, 32–39. (doi:10.1016/j.physbeh.2014.12.042).

[19] Bäckhed F, Fraser CM, Ringel Y, Sanders ME, Sartor RB, Sherman PM, Versalovic J, Young V, Finlay BB. 2012 Defining a healthy human gut microbiome: current concepts, future directions, and clinical applications. Cell host & microbe 12, 5, 611–622. (doi: 10.1016/j.chom.2012.10.012).

[20] The Human Microbiome Project Consortium. 2012 Structure, function and diversity of the healthy human microbiome. Nature 486, 7402, 207–214. (doi:10.1038/nature11234).

[21] Clavel T, Lagkouvardos I, Blaut M, Stecher B. 2016 The mouse gut microbiome revisited: From complex diversity to model ecosystems. International journal of medical microbiology: IJMM 306, 5, 316–327. (doi:10.1016/j.ijmm.2016.03.002).

[22] Chandler JA, Morgan Lang J, Bhatnagar S, Eisen JA, Kopp A. 2011 Bacterial communities of diverse drosophila species: Ecological context of a Host–Microbe model system. PLoS genetics 7, 9, e1002272. (doi:10.1371/journal.pgen.1002272).

[23] Wong CNA, Ng P, Douglas AE. 2011 Low-diversity bacterial community in the gut of the fruitfly drosophila melanogaster. Environmental microbiology 13, 7, 1889–1900. (doi: 10.1111/j.1462-2920.2011.02511.x).

[24] Koyle ML, Veloz M, Judd AM, Wong A, Newell PD, Douglas AE, Chaston JM. 2016 Rearing the fruit fly drosophila melanogaster under axenic and gnotobiotic conditions. Journal of visualized experiments: JoVE 113, e54219. (doi:10.3791/54219).

[25] Ali YO, Escala W, Ruan K, Zhai RG. 2011 Assaying locomotor, learning, and memory deficits in drosophila models of neurodegeneration. Journal of visualized experiments: JoVE, 49, e2504. (doi:10.3791/2504).

[26] Couzin ID. 2009 Collective cognition in animal groups. Trends in cognitive sciences 13, 1, 36–43. (doi:10.1016/j.tics.2008.10.002).

[27] Bates D, Mächler M, Bolker B, Walker S. 2015 Fitting linear mixed-effects models using lme4. Journal of Statistical Software 67, 1, 1–48. (doi:10.18637/jss.v067.i01).

[28] R Core Team. 2018 R: A Language and Environment for Statistical Computing. R Foundation for Statistical Computing, Vienna, Austria.

[29] Strausfeld NJ, Hirth F. 2013 Deep homology of arthropod central complex and vertebrate basal ganglia. Science 340, 6129, 157–161. (doi:10.1126/science.1231828).

[30] Apidianakis Y, Rahme LG. 2011 Drosophila melanogaster as a model for human intestinal infection and pathology. Disease models & mechanisms 4, 1, 21–30. (doi: 10.1242/dmm.003970).

[31] Qin J, Li R, Raes J, Arumugam M, Burgdorf KS, Manichanh C, Nielsen T, Pons N, Levenez F, Yamada T, et al.2010 A human gut microbial gene catalogue established by metagenomic sequencing. Nature 464, 7285, 59–65. (doi:10.1038/nature08821).

[32] Wang Z, Singhvi A, Kong P, Scott K. 2004 Taste representations in the drosophila brain. Cell 117, 7, 981–991. (doi:10.1016/j.cell.2004.06.011).

[33] Kim H, Kirkhart C, Scott K. 2017 Long-range projection neurons in the taste circuit of drosophila. eLife 6. (doi:10.7554/eLife.23386).

[34] Claridge-Chang A, Roorda RD, Vrontou E, Sjulson L, Li H, Hirsh J, Miesenböck G. 2009 Writing memories with light-addressable reinforcement circuitry. Cell 139, 2, 405–415. (doi:10.1016/j.cell.2009.08.034).

[35] Schwaerzel M, Monastirioti M, Scholz H, Friggi-Grelin F, Birman S, Heisenberg M. 2003 Dopamine and octopamine differentiate between aversive and appetitive olfactory memories in drosophila. Journal of Neuroscience 23, 33, 10495–10502.

[36] Schroll C, Riemensperger T, Bucher D, Ehmer J, Völler T, Erbguth K, Gerber B, Hendel T, Nagel G, Buchner E, et al. 2006 Light-induced activation of distinct modulatory neurons triggers appetitive or aversive learning in drosophila larvae. Current biology: CB 16, 17, 1741–1747. (doi:10.1016/j.cub.2006.07.023).

[37] Kim YC, Lee HG, Han KA. 2007 D1 dopamine receptor dDA1 is required in the mushroom body neurons for aversive and appetitive learning in drosophila. The Journal of neuroscience: the official journal of the Society for Neuroscience 27, 29, 7640–7647. (doi: 10.1523/JNEUROSCI.1167-07.2007).

[38] Liu X, Davis RL. 2009 The GABAergic anterior paired lateral neuron suppresses and is suppressed by olfactory learning. Nature Neuroscience 12, 1, 53–59. (doi:10.1038/nn.2235).

[39] Pavlowsky A, Schor J, Plaçais PY, Preat T. 2018 A GABAergic feedback shapes dopaminergic input on the drosophila mushroom body to promote appetitive Long-Term memory. Current biology: CB 28, 11, 1783–1793.e4. (doi:10.1016/j.cub.2018.04.040).

[40] Monier M, Nbel S, Danchin E, Isabel G. 2019 Dopamine and Serotonin Are Both Required for Mate-Copying in Drosophila melanogaster. Frontiers in Behavioral Neuroscience 12. (doi:10.3389/fnbeh.2018.00334).

[41] Wise RA, Rompre PP. 1989 Brain dopamine and reward. Annual review of psychology 40, 191–225. (doi:10.1146/annurev.ps.40.020189.001203).

[42] Schultz W, Dayan P, Montague PR. 1997 A neural substrate of prediction and reward. Science 275, 5306, 1593–1599. (doi:10.1126/science.275.5306.1593).

[43] Myhrer T. 2003 Neurotransmitter systems involved in learning and memory in the rat: a meta-analysis based on studies of four behavioral tasks. Brain Research Reviews 41, 2-3, 268–287. (doi:10.1016/S0165-0173(02)00268-0).

[44] Foster JA. 2016 Chapter three - gut microbiome and behavior: Focus on neuroimmune interactions. In: Cryan JF, Clarke G (eds.), International Review of Neurobiology, vol. 131, pp. 49–65. Academic Press. (doi:10.1016/bs.irn.2016.07.005).

[45] Liu YW, Liu WH, Wu CC, Juan YC, Wu YC, Tsai HP, Wang S, Tsai YC. 2016 Psychotropic effects of Lactobacillus plantarum PS128 in early life-stressed and nave adult mice. Brain Research 1631, 1–12. (doi:10.1016/j.brainres.2015.11.018).

[46] Desbonnet L, Garrett L, Clarke G, Bienenstock J, Dinan TG. 2008 The probiotic bifidobacteria infantis: An assessment of potential antidepressant properties in the rat. Journal of psychiatric research 43, 2, 164–174. (doi:10.1016/j.jpsychires.2008.03.009).

[47] Collins SM, Kassam Z, Bercik P. 2013 The adoptive transfer of behavioral phenotype via the intestinal microbiota: experimental evidence and clinical implications. Current opinion in microbiology 16, 3, 240–245. (doi:10.1016/j.mib.2013.06.004).

[48] Hanstock T, Clayton E, Li K, Mallet P. 2004 Anxiety and aggression associated with the fermentation of carbohydrates in the hindgut of rats. Physiology & Behavior 82, 2-3, 357–368. (doi:10.1016/j.physbeh.2004.04.002).

[49] Hanstock T, Mallet P, Clayton E. 2010 Increased plasma d-lactic acid associated with impaired memory in rats. Physiology & Behavior 101, 5, 653–659. (doi: 10.1016/j.physbeh.2010.09.018).

[50] Unger MM, Spiegel J, Dillmann KU, Grundmann D, Philippeit H, Brmann J, Fabender K, Schwiertz A, Schfer KH. 2016 Short chain fatty acids and gut microbiota differ between patients with Parkinson’s disease and age-matched controls. Parkinsonism & Related Disorders 32, 66–72. (doi:10.1016/j.parkreldis.2016.08.019).

[51] Cho T, Lee C, Lee N, Hong YR, Koo J. 2019 Small-chain fatty acid activates astrocytic odorant receptor Olfr920. Biochemical and Biophysical Research Communications 510, 3, 383–387. (doi:10.1016/j.bbrc.2019.01.106).

[52] Hang S, Purdy AE, Robins WP, Wang Z, Mandal M, Chang S, Mekalanos JJ, Watnick PI. 2014 The acetate switch of an intestinal pathogen disrupts host insulin signaling and lipid metabolism. Cell host & microbe 16, 5, 592–604. (doi:10.1016/j.chom.2014.10.006).

[53] Wong ACN, Vanhove AS, Watnick PI. 2016 The interplay between intestinal bacteria and host metabolism in health and disease: lessons from drosophila melanogaster. Disease models & mechanisms 9, 3, 271–281. (doi:10.1242/dmm.023408).

[54] Kamareddine L, Robins WP, Berkey CD, Mekalanos JJ, Watnick PI. 2018 The drosophila immune deficiency pathway modulates enteroendocrine function and host metabolism. Cell metabolism 28, 3, 449–462.e5. (doi:10.1016/j.cmet.2018.05.026).

[55] Frost G, Sleeth ML, Sahuri-Arisoylu M, Lizarbe B, Cerdan S, Brody L, Anastasovska J, Ghourab S, Hankir M, Zhang S, et al. 2014 The short-chain fatty acid acetate reduces appetite via a central homeostatic mechanism. Nature Communications 5, 1. (doi: 10.1038/ncomms4611).

[56] Garai G, Dueas M, Irastorza A, Moreno-Arribas M. 2007 Biogenic amine production by lactic acid bacteria isolated from cider. Letters in Applied Microbiology 45, 5, 473–478. (doi:10.1111/j.1472-765X.2007.02207.x).

